# Temperature and O_2_, but not CO_2_, interact to affect aerobic performance of European sea bass (Dicentrarchus labrax)

**DOI:** 10.1101/2021.03.12.435078

**Authors:** Daniel W. Montgomery, Stephen D. Simpson, William Davison, Harriet R. Goodrich, Georg H. Engelhard, Silvana N.R. Birchenough, Rod W. Wilson

## Abstract

Climate change causes warming, decreased O_2_, and increased CO_2_ in marine systems and responses of organisms will depend on interactive effects between these factors. We provide the first experimental assessment of the interactive effects of warming (14 to 22°C), reduced O_2_ (∼3 – 21 kPa O_2_), and increased CO_2_ (∼400 or ∼1000 µatm ambient CO_2_) on four indicators of aerobic performance (standard metabolic rate, SMR, maximum metabolic rate, MMR, aerobic scope, and hypoxia tolerance, O_2crit_), blood chemistry, and O_2_ transport (P_50_) of a marine fish, the European sea bass (Dicentrarchus labrax). Warming increased SMR and O_2crit_ (i.e. reduced hypoxia tolerance) as well as MMR in normoxia but there was an interactive effect with O_2_ so that hypoxia caused larger reductions in MMR and aerobic scope at higher temperatures. Increasing CO_2_ had minimal effects on SMR, MMR and O_2crit_ and did not show interactive effects with temperature or O_2_ for any measured variables. Aerobic performance was not linked to changes in blood chemistry or P_50_. Despite lack of effects of CO_2_ on aerobic performance, increased CO_2_ induced 30% mortality of fish exercised in low O_2_ at 22°C indicating important threshold effects independent of aerobic performance. Overall, our results show temperature and O_2_, but not CO_2_, interact to affect aerobic performance of sea bass, disagreeing with predictions of the oxygen- and capacity-limited thermal tolerance hypothesis.

## 1. Introduction

Atmospheric CO_2_ levels are increasing and could reach ∼1000 µatm by the end of the century (IPCC, 2014). Rising atmospheric greenhouse gases increase ocean temperatures (Bopp et al., 2013), which reduces oceanic O_2_ content and exacerbates the frequency and severity of hypoxic (low O_2_) events (Breitburg et al., 2018; Diaz & Rosenberg, 2008). As atmospheric CO_2_ levels continue to rise so too does the concentration of CO_2_ in marine systems (Caldeira & Wickett, 2003). Therefore, responses of marine organisms to climate change will be a result of simultaneous changes in temperature, O_2_, and CO_2_.

Changes in temperature, O_2_, and CO_2_ can individually impact physiological performance of fish. Concern has been raised that interactions between these three may occur in a non-linear manner, so that their combined impact cannot be predicted from responses to an individual variable (Côté et al., 2016; Crain et al., 2008; McBryan et al., 2013; Todgham & Stillman, 2013). As such, we need to understand how temperature, O_2_, and CO_2_ interact to affect the physiology of fish to enable accurate predictions of how climate change will influence fish species (Hollowed et al., 2013; Pörtner & Peck, 2010; Wernberg et al., 2012). One approach is to examine how these environmental factors affect the fish’s range of aerobic metabolism (aerobic scope), defined as the difference between an animal’s maximum rate of O_2_ consumption (maximum metabolic rate, MMR) and the minimum rate needed to meet basal energy demands (standard metabolic rate, SMR) (Fry, 1971).

It has been proposed that aerobic scope provides a single metric of whole-animal performance in a particular environment. Therefore, aerobic scope can be directly linked to processes such as growth and reproduction and in turn overall organism fitness, through the concept of oxygen- and capacity-limited thermal tolerance, OCLTT (Pörtner, 2012; Pörtner et al., 2017). The OCLTT hypothesis assumes that an organism’s overall fitness is maximised at an optimal temperature at which aerobic scope peaks. As such, anything that reduces aerobic scope will also diminish fitness, potentially changing the distribution of populations (Pörtner & Farrell, 2008). Hypoxia affects aerobic scope by limiting environmental O_2_ availability and, therefore, the maximum O_2_ uptake rate possible by fish. Increased CO_2_ has been proposed to affect aerobic scope both by increasing the SMR of fish (e.g. through increased cost of acid-base regulation) and by decreasing MMR (potentially because subsequent changes in internal acid-base chemistry can reduce O_2_ transport capacity of the blood or impair tissue functioning). The OCLTT hypothesis therefore predicts that reduced O_2_ and increased CO_2_ will interact to reduce aerobic scope across the thermal performance curve of a species. This would result in a lower optimal temperature (where peak aerobic scope occurs) and reduced thermal tolerance. While this concept has been successfully used to explain changes in habitat suitability and population distributions of some fish species (Cucco et al., 2012; Del Raye & Weng, 2015), the assertion that it represents a universal framework to predict effects of climate change on all fish populations (Farrell, 2016) has been challenged (Jutfelt et al., 2018; Lefevre, 2016).

The proposal of the OCLTT hypothesis has led to numerous studies examining how O_2_ or CO_2_ interact with temperature to affect aerobic performance (for examples see Chabot and Claireaux, 2008; Rummer et al., 2013; Grans et al., 2014). However, to date no experimental work has sought to investigate how combined changes in all three factors (temperature, O_2_, and CO_2_) interact to affect aerobic performance in fish. Combining all three environmental variables is vital to accurately assess potential interactive effects for three reasons. Firstly, meta-analysis of multi-factor studies indicates that the prevalence of non-linear effects doubles when moving from studies that combine two factors to three factors (Crain et al., 2008). Secondly, the role of CO_2_ as a limiting factor of aerobic scope was originally suggested to occur primarily in combination with hypoxia (Fry, 1971). Thirdly, low O_2_ conditions in the environment always co-occur with increased CO_2_ (Melzner et al., 2013). As such, experiments investigating effects of O_2_ or CO_2_ and temperature on aerobic performance may not accurately reflect interactive effects caused by all three environmental factors.

In this study we investigated how temperature, O_2_, and CO_2_ interact to affect aerobic performance of European sea bass (Dicentrarchus labrax), a species showing recent northward range expansions thought to be related to warming (Pawson et al., 2007). Two separate populations exist (Souche et al., 2015) and, although the physiological responses of this species to environmental change have been regularly examined, to date only one study has used fish from the Atlantic population. As such, our experiment had three aims:

i. to assess how aerobic performance of sea bass from the Atlantic population will respond to predicted future environmental changes;
ii. to determine whether combinations of hypoxia and increased CO_2_ interact with temperature to affect aerobic scope, as predicted by the OCLTT hypothesis;
iii. to determine whether changes in aerobic performance are linked to blood chemistry and O_2_ transport capacity.

## 2. Results

### 2.1. Standard metabolic rate and hypoxia tolerance

The best supported model for SMR included both temperature and CO_2_ as fixed effects (Linear Mixed-Effects Model, marginal R^2^ = 0.70, conditional R^2^ = 0.70, Table S6). Standard metabolic rate approximately doubled between 14 and 22°C, exhibiting a Q_10_ temperature coefficient of 2.09 (Figure 1). The best model indicated that CO_2_ had a small effect on SMR - reducing SMR by 7.4 mgO_2_ kg^-1^ h^-1^, ∼10 % of the smallest SMR, (95 % CI = -1.65 to 16.43 mgO_2_ kg^-1^ h^-1^) across all temperatures at ∼1000 µatm CO_2_ (Figure 1). However, the model including temperature but not CO_2_ had a ΔAICc <2 indicating that including the effect of CO_2_ in the best model did not lead to a large improvement in model fit (Table S5). There was no evidence of an interactive effect between increasing temperatures and increasing CO_2_.

**Figure 1:**
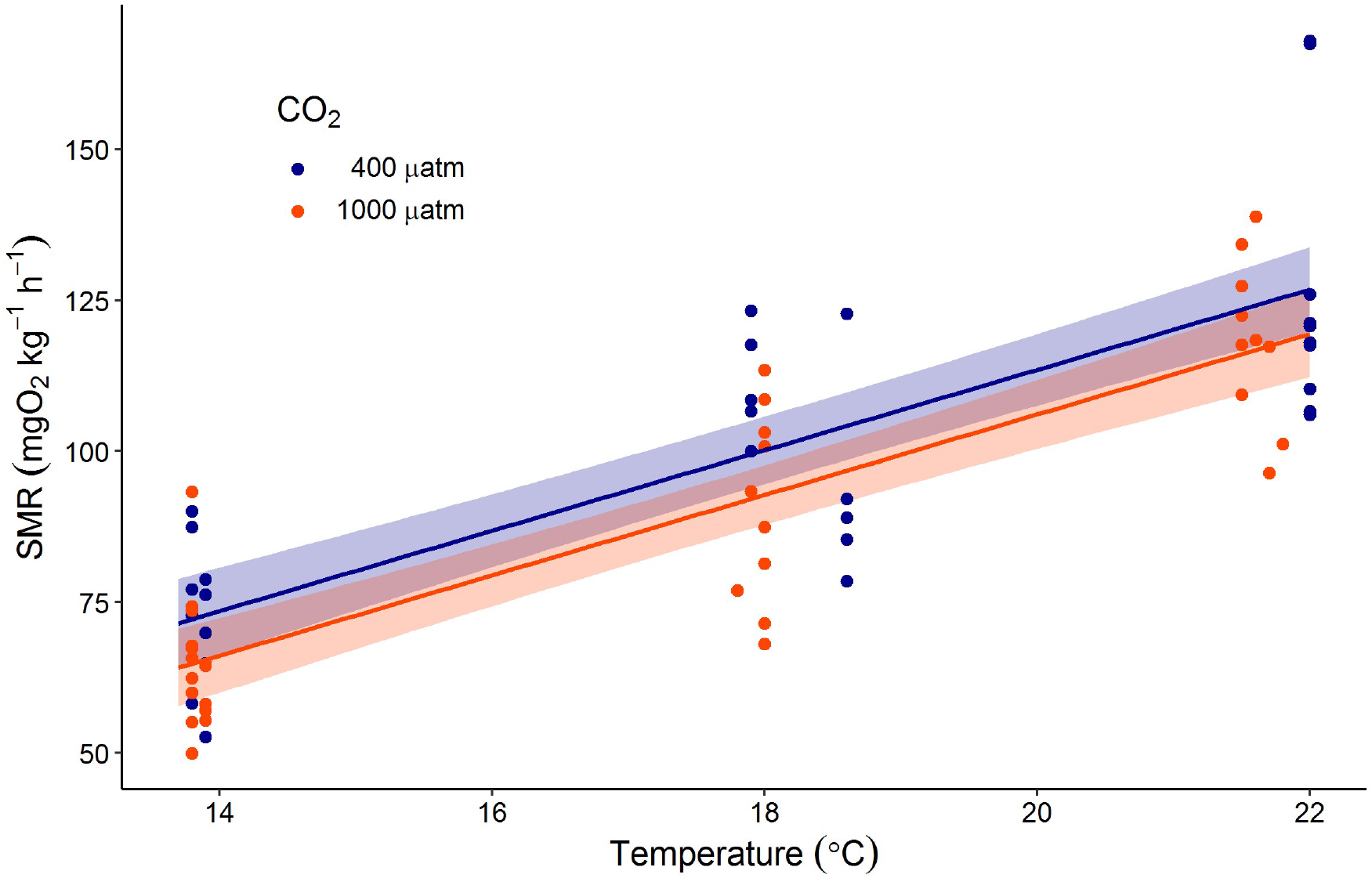
Impact of temperature and CO_2_ on standard metabolic rates (SMR) of juvenile sea bass. The best supported model (marginal R^2^ = 0.70, conditional R^2^ = 0.70) to explain variation in SMR included temperature and CO_2_ as explanatory variables but not their interaction. Points represent calculated SMR for individual fish, lines represented predicted SMR at two CO_2_ levels (blue = present day ∼400 µatm CO_2_, orange = end of century ∼ 1000 µatm CO_2_) from the best supported model, and shaded areas represent bootstrapped 95 % CI of predictions (n = 1000).

The best supported model of O_2crit_ included the effects of temperature, CO_2_ and SMR but no interactions (Linear Mixed-Effects Model, marginal R^2^ =0.72, conditional R^2^ = 0.77, Table S7). A doubling in SMR from 60 to 120 mgO_2_ kg^-1^ h^-1^ is predicted to increase O_2crit_ by 2.16 kPa O_2_ (95 % CI = 1.66 to 2.66 kPa O_2_). Independent of their effects on SMR, both temperature and CO_2_ were included in the best model of O_2crit_. The effect of temperature meant that for a given value of SMR O_2crit_ would reduce as temperature increased. For instance, a fish at 14 °C is predicted to have an O_2crit_ 0.50 kPa O_2_ (95 % CI = 0.13 to 0.87 kPa O_2_) higher than a fish with the same SMR at 18 °C (Figure 2A). Combined the effects of SMR and temperature mean warming from 14 to 22 °C will increase O_2crit_ by 0.91 kPa O_2_ (95 % CI = 0.53 to 1.29 kPa O_2_, Figure 2B). The additional effect of CO_2_ is predicted to increase O_2crit_. Despite this, when taking into account the effect of CO_2_ on SMR from the best model of SMR (Figure 1), and the effect of SMR and temperature on O_2crit_, the resulting effect of CO_2_ causes an increase in O_2crit_ of only 0.04 kPa O_2_ (95 % CI = -0.32 to 0.40 kPa O_2_, Figure 2B).

**Figure 2:**
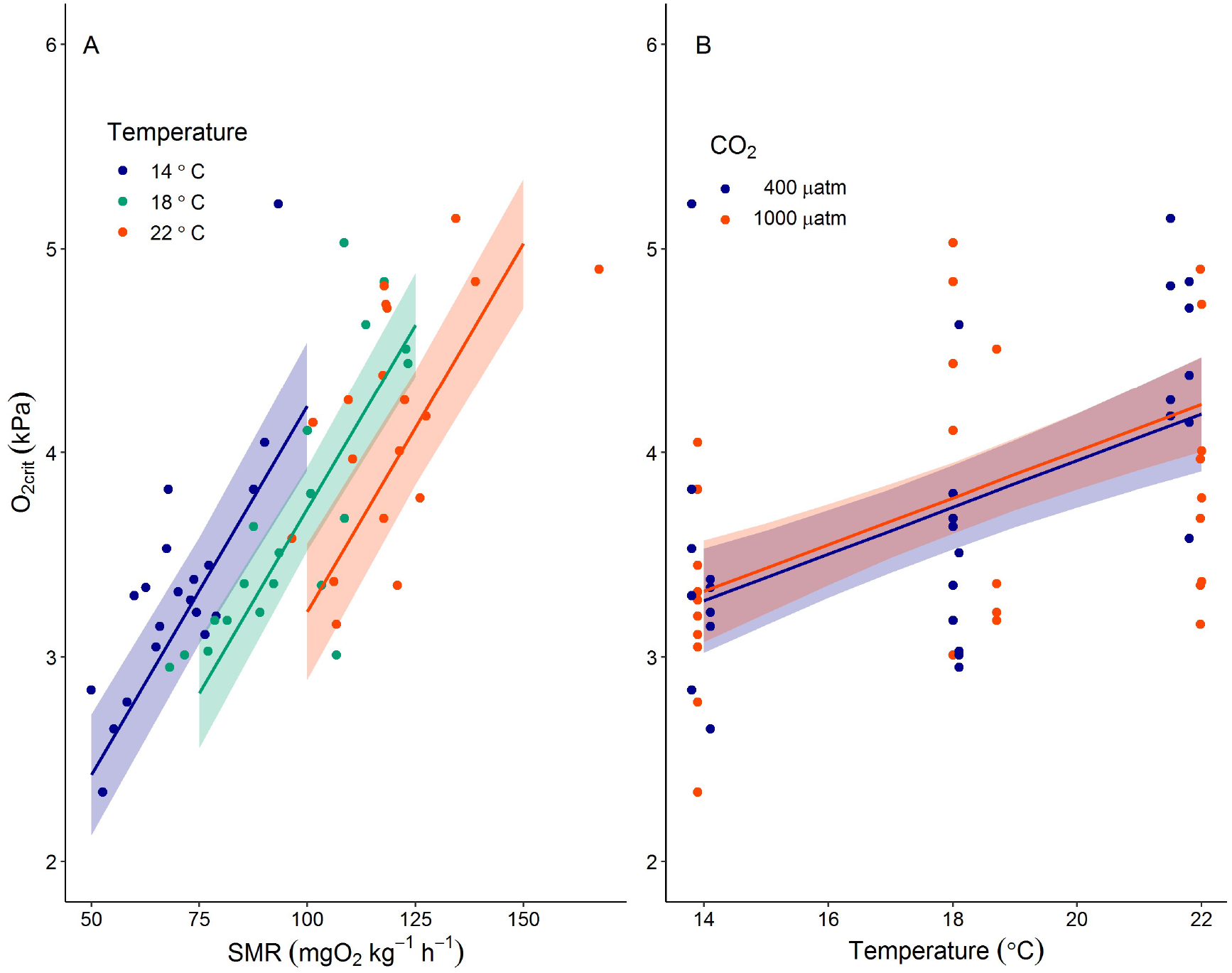
Combined impacts of SMR, temperature and CO_2_ on hypoxia tolerance (O_2crit_) of juvenile European sea bass. A. The best supported model (marginal R^2^ = 0.72, conditional R^2^ = 0.77) predicted a positive effect of increasing SMR on O_2crit_ with the effect of increased temperature resulting in lower O_2crit_ for a given value of SMR. B. Combined effects of SMR and temperature result in an increase in O_2crit_ between 14 °C and 22 °C. The predicted positive effect of increased CO_2_ on O_2crit_ is small compared to changes in SMR and temperature. Points represent calculated O_2crit_ for individual fish, lines represented predicted O_2crit_ from the best supported model, and shaded areas represent bootstrapped 95 % CI of predictions (n = 1000).

### 2.2. Maximum metabolic rate and aerobic scope

Maximum metabolic rate of sea bass was affected by temperature, O_2_, and CO_2_ level. Warming from 14 to 22 °C, in normoxia combined with normocapnia, caused a 50 % increase in MMR from 293.5 ± 10.4 to 435.9 ± 18.9 mgO_2_ kg^-1^ h^-1^, with a Q_10_ of 1.64. In high CO_2_ the temperature effect on MMR was very similar (Q_10_ = 1.60). The best supported model predicted that O_2_ had a non-linear quadratic effect on MMR, so that a given reduction in O_2_ caused a larger reduction in MMR at lower O_2_ levels, as well as interactive effects between O_2_ and temperature (Figure 3; Linear Mixed Model, marginal R^2^ = 0.91, conditional R^2^ = 0.96, Table S8). For example a 5 kPa reduction in O_2_ at 18 °C (in normal CO_2_) from air saturated levels (∼ 20 to ∼15 kPa O_2_) results in a predicted decrease in MMR of 8.0 mgO_2_ kg^-1^ h^-1^ (95 % CI = -7.2 to 23.2 mgO_2_ kg^-1^ h^-1^) whereas the same reduction in O_2_ from 10 to 5 kPa resulted in a 15-fold larger predicted decrease in MMR of 123.8 mgO_2_ kg^-1^ h^-1^ (95 % CI = 115.1 to 132.5 mgO_2_ kg^-1^ h^-1^). In addition, the non-linear O_2_ effect interacted with the temperature effect so that the same reduction in O_2_ caused a larger reduction in MMR as temperature increased (Figure 3A & B). This was particularly noticeable between 15 and 10 kPa O_2_ where fish at 22 °C (in normal CO_2_) exhibited a predicted decline in MMR that was 60 % larger than fish at 14 °C, 80.9 mgO_2_ kg^-1^ h^-1^ (95 % CI = 63.2 to 98.6 mgO_2_ kg^-1^ h^-1^) versus 50.8 mgO_2_ kg^-1^ h^-1^ (95 % CI = 35.9 to 65.7 mgO_2_ kg^-1^ h^-1^). Finally, environmental CO_2_ level had a small, negative effect independent of interactions between temperature and O_2_. As a result, an increase in CO_2_ of 1000 µatm is predicted to reduce MMR by 8.5 mgO_2_ kg^-1^ h^-1^ (95 % CI = -18.2 to 35.3 mgO_2_ kg^-1^ h^-1^) irrespective of temperature and O_2_ (Figure 3C, D, & E).

**Figure 3:**
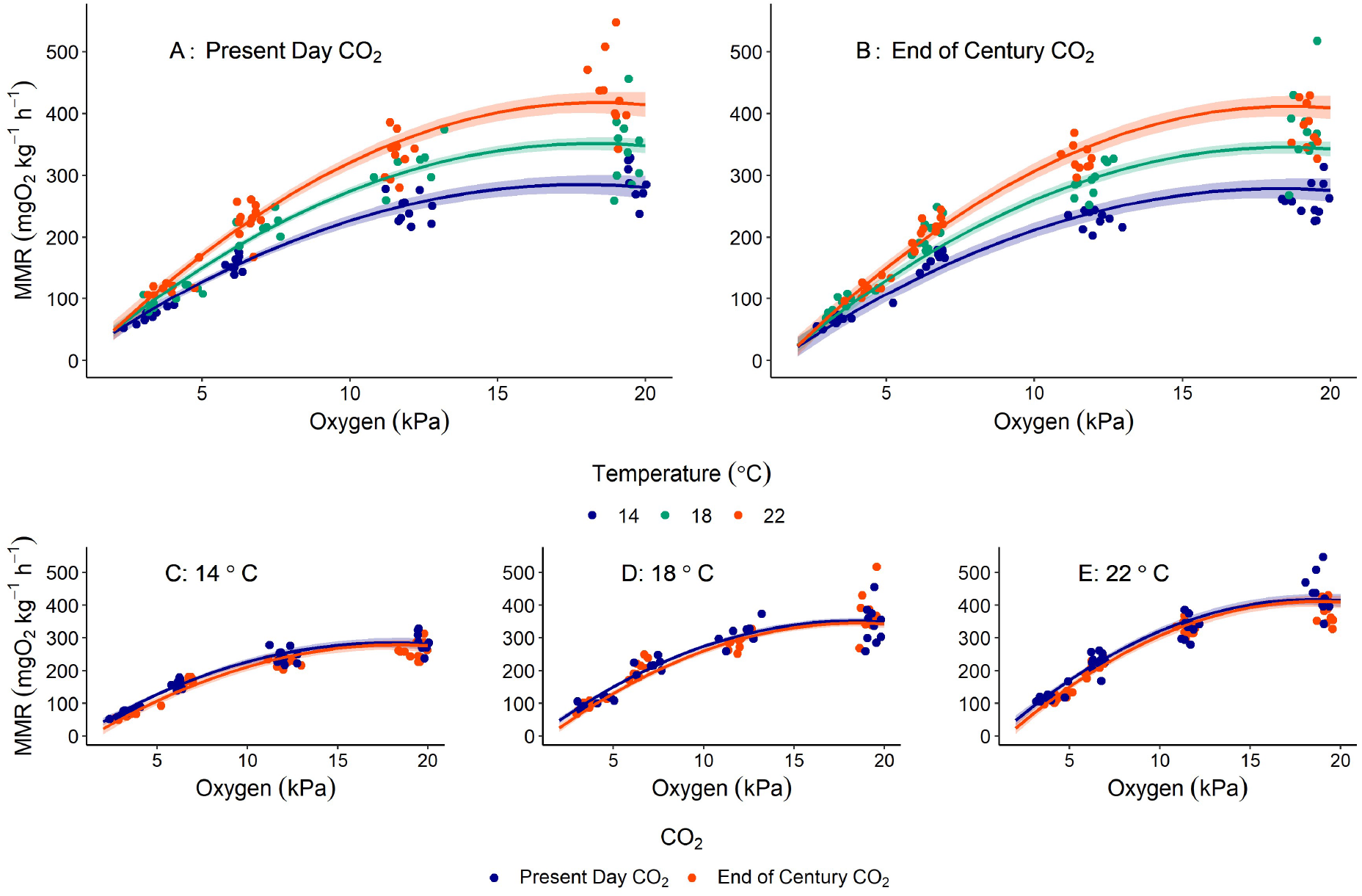
Effects of combinations of temperature, O_2_, and CO_2_ on the maximum metabolic rate (MMR) of European sea bass. There was a synergistic interactive effect of temperature and O_2_ on MMR for sea bass exposed to both A. present day CO_2_ conditions (present day CO_2_ = 400 µatm at ∼20 kPa O_2_) and B. end of century CO_2_ conditions (end of century CO_2_ = 1000 µatm at ∼ 20 kPa O_2_). The additive effects of reduced O_2_ and increased CO_2_ levels are displayed for sea bass at A. 14 °C B. 18 °C and C. 22 °C. Points represent calculated MMR for individual fish, lines represent predicted MMR from the best supported model, and shaded areas represent bootstrapped 95 % confidence intervals (n = 1000).

We predicted the impacts of temperature, O_2_, and CO_2_ on aerobic scope from the best supported models fitted to measurements of SMR and MMR (Figure 4). At normoxia (∼20 kPa O_2_) aerobic scope is predicted to increase by 80.2 mgO_2_ kg^-1^ h^-1^ (95 % CI = 24.8 to 135.7 mgO_2_ kg^-1^ h^-1^) as temperature increases from 14 to 22 °C independent of changes in CO_2_ level (Figure 4A & B). Interactive effects between temperature and O_2_ on MMR are reflected in predictions of aerobic scope (Figure 4A &B). As increasing CO_2_ has the same direction of effect on both SMR and MMR the impact of CO_2_ on aerobic scope is minimal (Figure 4C, D, & E).

**Figure 4:**
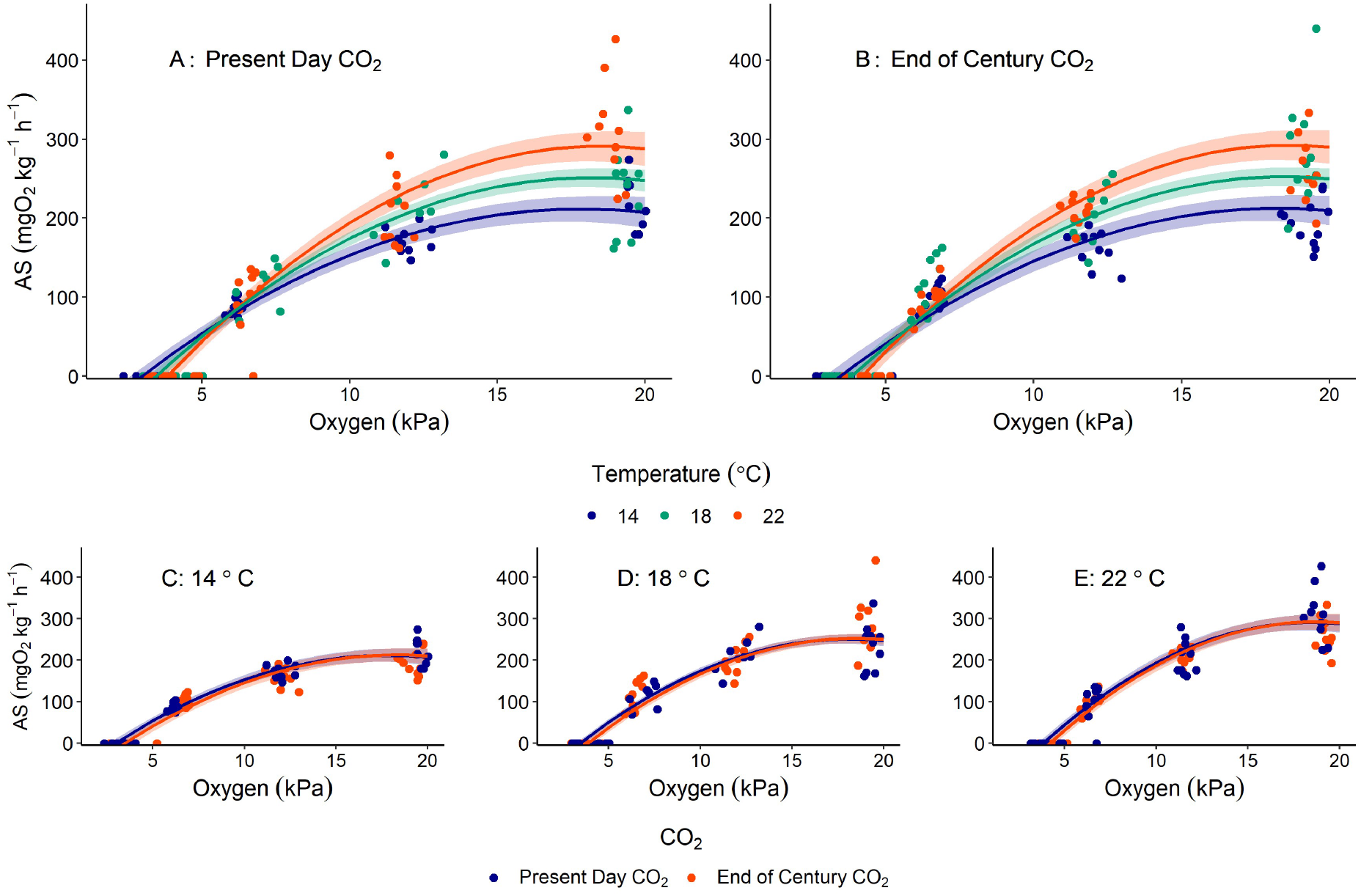
Effects of combinations of temperature, O_2_, and CO_2_ on the aerobic scope (AS) of European sea bass. There was a non-linear interactive effect of temperature and O_2_ on aerobic scope for sea bass exposed to both A. present day CO_2_ conditions (present day CO_2_ = 400 µatm at ∼20 kPa O_2_) and B. end of century CO_2_ conditions (end of century CO_2_ = 1000 µatm at ∼ 20 kPa O_2_). The additive effects of reduced O_2_ and increased CO_2_ levels are displayed for sea bass at A. 14 °C B. 18 °C and C. 22 °C. Points represent aerobic scope of individual fish derived from calculated RMR and MMR of that individual, lines represent predicted aerobic scope calculated by subtracting model predictions of RMR and MMR and shaded areas represent 95 % confidence intervals calculated from bootstrapped standard errors of predicted RMR and MMR (n = 1000).

### 2.3. Blood chemistry & Hb-O_2_ affinity

Increasing ambient CO_2_ from ∼400 to ∼1000 µatm increased plasma pCO_2_ (Two-way ANOVA, F = 15.84, df = 1, p < 0.001) whereas warming from 14 to 22 °C did not (F = 0.772, df = 2, p = 0.468), and there was no interactive effect noted (F = 2.067, df = 2, p = 0.138). Despite the significant overall effect of increased CO_2_ on plasma pCO_2_ pairwise comparisons (Figure 5) only revealed a significant increase in plasma pCO_2_ in bass at 18 °C exposed to ∼1000 µatm (0.53 ± 0.05 kPa pCO_2_) compared to ambient conditions (0.29 ± 0.05 kPa pCO_2_) (Pairwise comparisons of least square means, t = 3.688, df = 1, p = 0.008).

**Figure 5:**
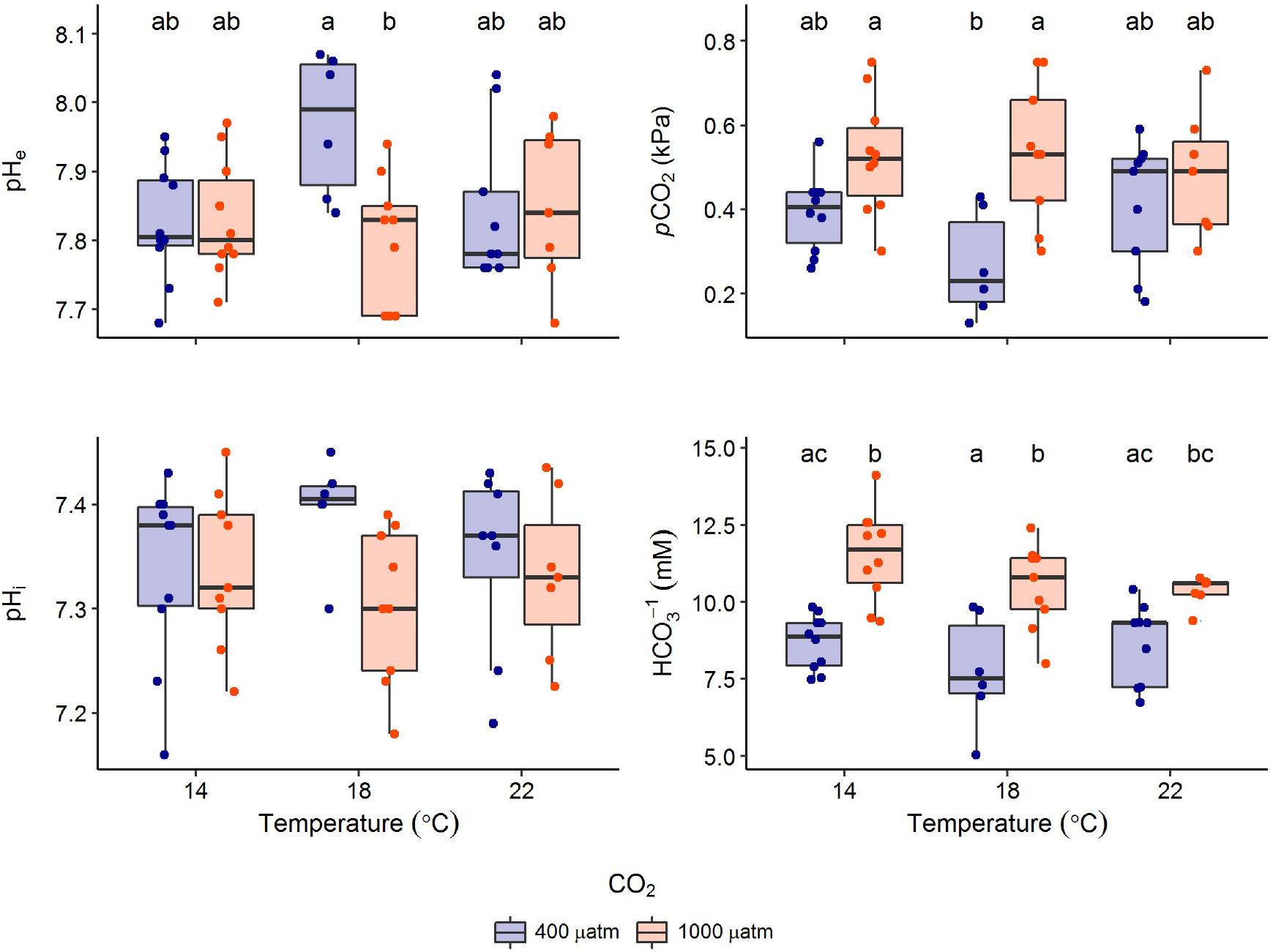
Impact of temperature and CO_2_ on blood acid-base characteristics of European sea bass. Significant differences in pairwise comparisons of least square means between treatments are indicated by different letters (a,b,c). No significant differences between any treatments were noted for measurements of pH_i_.

To compensate for increased plasma pCO_2_ in ∼1000 µatm CO_2_ treatments sea bass accumulated ∼3 mM extra HCO_3_^-^ (Two-way ANOVA, F = 44.34, df = 1, p << 0.001) (Figure 5), compared to fish in ambient CO_2_ conditions at 14 and 18 °C. Fish exposed to ∼1000 µatm CO_2_ at 22 °C showed a non-significant increase in HCO_3_^-^ of just under 2 mM (95 % CI = 0.60 to 2.84 mM) when compared to fish at ∼400 µatm CO_2_ (t = 2.66, df = 1, p = 0.103). There was no temperature effect on plasma HCO_3_^-^ (F = 2.538, df = 2, p = 0.0903) and no interactive effect between temperature and CO_2_ (F = 0.969, df = 2, p = 0.387).

As a result of compensatory accumulation of HCO_3_^-^, blood extracellular pH (pH_e_) was regulated in response to high CO_2_ (F = 3.56, df = 1, p = 0.066) and sea bass did not show significant effects of temperature on pH_e_ (Two way ANOVA, F = 1.425, df = 2, p = 0.251). However, there was a significant interactive effect between temperature and CO_2_ (F = 3.952, df = 2, p = 0.026). This interactive effect was caused by a significant reduction in pH_e_ in sea bass exposed to ∼1000 µatm CO_2_ at 18 °C (7.80 ± 0.03) when compared to fish at ambient CO_2_ levels (7.97 ± 0.04) (pairwise comparisons of least square means, t = 3.242, df = 1, p = 0.026). We did not find significant differences between pH_e_ across all other treatment groups. Additionally, intracellular pH of red blood cells (pH_i_) showed no significant differences between all treatments (Kruskal-Wallis test, χ^2^ = 6.79, df = 5, p = 0.237) (Figure 5).

There were no significant differences in plasma lactate levels across all treatments (Kruskall-Wallis, χ^2^ = 6.40, df = 5, p = 0.269), with mean lactate for all fish of 0.42 ± 0.06 mM (± S.E.). Plasma glucose levels (mean for all fish of 4.30 ± 0.14 mM, ± S.E.) were not significantly affected by temperature (Two-way ANOVA, F = 0.864, df = 2, p = 0.429), CO_2_ (F = 0.0138, df = 1, p = 0.907) or the interaction between temperature and CO_2_ (F = 1.052, df = 2, p = 0.358).

Our measurements of blood O_2_ transport showed no consistent impacts of temperature or CO_2_. Haemoglobin-O_2_ binding affinity (P_50_) (Kruskal-Wallis, χ^2^ = 6.503, df = 4, p = 0.165) and Hills’ number (One-way ANOVA, F = 0.878, df = 4, p = 0.487) were not significantly different between any treatments (Figure S1). Haematocrit (Hct) was not affected by temperature (Two-way ANOVA, F = 0.832, df = 2, p = 0.442) or CO_2_ (F = 2.945, df = 1, p = 0.093) and no interactive effects were evident (F = 0.069, df = 2, p = 0.934). Haemoglobin (Hb) levels were affected by temperature (Two-way ANOVA, F = 7.094, df = 2, p = 0.002) with fish sampled at 22 °C having Hb levels 0.20 and 0.24 mM higher than fish sampled at 18 and 14 °C respectively. Haemoglobin levels were not affected by CO_2_ (F = 0.690, df = 1, p = 0.411) and there were no interactive effects (F = 3.067, df = 2, p = 0.057) (Figure S2).

### 2.4. Mortality

Three fish (30 %) died ∼1 hour post-chase after exposure to ∼6 kPa O_2_ at 22 °C in the high CO_2_ treatment. No fish died post-chase in any other treatment combinations.

## 3. Discussion

Our results demonstrate, for the first time, the interactive effects of temperature, O_2_, and CO_2_ on aerobic performance of an active predatory marine fish. We show that temperature and O_2_ have a non-linear interactive effect on aerobic performance of European sea bass but CO_2_’s impact is minor and independent. Both SMR and MMR increased with temperature from 14 to 22 °C, but changes in MMR were greater leading to positive effects on absolute aerobic scope. These results suggest that European sea bass populations in the North-East Atlantic (typical temperatures are < 22 °C) could physiologically benefit from global warming. However, hypoxia tolerance reduced at higher temperatures, and hypoxia reduced MMR and aerobic scope by a greater amount at high temperatures, indicating that in warmer waters European sea bass will be more susceptible to hypoxia.

We confirm similar findings from previous research investigating effects of temperature and O_2_ on European sea bass. For example, the increase in SMR with temperature we observed closely follows results from aquaculture sourced European sea bass (Claireaux & Lagardère, 1999). That is the Q_10_ temperature coefficient dropped from 2.33 between 14 and 18 °C to 1.82 between 18 and 22 °C. This drop in Q_10_ as temperature increases provides further evidence that SMR does not increase exponentially after chronic vs acute exposure (Sandblom et al., 2014; Schulte et al., 2011). Responses of MMR to temperature showed similar Q_10_’s indicating a linear increase in MMR between 14 and 22 °C. This also closely corresponds with observations from Claireaux and Lagardère (1999) who found that MMR increased approximately linearly between 14 and 22 °C, before peaking between 22 and 24 °C and declining at higher temperatures. Further work with sea bass from Mediterranean stock has shown that specific growth rate and feed conversion efficiency peak at ∼25 °C and decline at higher temperatures (Person-Le Ruyet et al., 2004), similar to changes in metabolic scope shown by Claireaux and Lagardère (1999). Finally, aquaculture-produced sea bass display an Arrhenius break point of heart rate at ∼21.5 °C and developed arrhythmia at ∼26 °C (Crespel et al., 2019). The consistency in temperature of peak performance in sea bass from distinct sub-populations with vastly different environmental experiences supports the idea that fish face ceilings to physiological performance in the face of environmental change (Sandblom et al., 2016). Despite these similarities with previous research, SMR of fish in our study was higher at a given temperature than for fish from Mediterranean stocks (Claireaux & Lagardère, 1999). This may support the theory of metabolic cold adaptation, that basal energy demand in fish from warmer environments will be lower than in fish from cold environments when measured at the same temperature (Krogh, 1916). This has recently been supported by evidence from wild populations of three-spine stickleback (Gasterosteus aculeatus) (Pilakouta et al., 2020).

Declining O_2_ caused a decrease in MMR along a limiting oxygen curve, similar to that seen previously in sea bass (Claireaux & Lagardère, 1999; Lagardère et al., 1998) and other fish species (Chabot & Claireaux, 2008; Claireaux et al., 2000; Lefrancois & Claireaux, 2003; Mallekh & Lagardère, 2002). This result questions recent predictions made by Seibel and Deutsch (2020) that MMR of fish should decrease linearly from ∼21 kPa O_2_ to O_2crit_ values. The curved, rather than linear, response of MMR we observed may occur as a result of compensatory mechanisms (e.g. increased ventilation, cardiac output, gill lamellar perfusion and surface area, and haematocrit) to maintain O_2_ delivery (Farrell & Richards, 2009). While these adjustments may limit reductions in MMR during mild to moderate hypoxia they may reach their performance limits as O_2_ approaches critical levels, resulting in a steeper decline in MMR. For example, in moderate hypoxia rainbow trout (Oncorhynchus mykiss) increase cardiac output via increased stroke volume but in severe hypoxia cardiac output cannot be increased further leading to bradycardia (Sandblom & Axelsson, 2005).

Temperature and O_2_ interacted to affect metabolism of sea bass so that impacts of hypoxia on MMR increased with temperature (Figure 3). This result supports previous research by Claireaux & Lagardère (1999). However, sea bass in our study displayed higher MMR at similar temperature and pO_2_ when compared to bass from Claireaux and Lagardère (1999). In addition, the O_2crit_ (the point at which SMR = MMR) of our sea bass increased between 14 °C and 22 °C (Figure 2) whereas results from previous work suggested that O_2crit_ increases or remains constant across this temperature range (Claireaux & Lagardère, 1999). The reduction in hypoxia tolerance of sea bass with warming was primarily a result of strong positive correlation between O_2crit_ and SMR (Figure 1). This relationship has been shown for numerous fish species, and most recently in work with black sea bass (Centropristus striata) (Slesinger et al., 2019). However, our results also indicate that temperature had a secondary effect which resulted in lower O_2crit_ at higher temperatures for a given SMR. This suggests that temperature affects O_2crit_ via another mechanism (or mechanisms) independent of SMR. This is unlikely to be related to O_2_ transport capacity of the blood as there were few consistent effects of temperature on Hct, Hb or P_50_ of sea bass (Figure 5, S1 and S2). It has previously been observed that improved hypoxia tolerance of fish after acclimation to increased temperature correlates to increased gill lamellar surface area (McBryan et al., 2016) or changes to heart structure (Anttila et al., 2015). Gomez Isaza et al. (2021) demonstrated that both of these cardiorespiratory responses can improve O_2_ supply capacity. Combined, these results suggest that thermal acclimation can cause structural changes to the gills and the heart, improving performance in low O_2_ conditions and mitigating the negative effect of temperature induced increases in SMR on hypoxia tolerance.

The independent effect of rising CO_2_ reduced MMR. Previous research has not shown consistent effects of CO_2_ on MMR or SMR of fish species, with the majority showing no effects of CO_2_ (Lefevre, 2016, 2019). Interestingly the most recent research with European sea bass found that long term exposure to elevated CO_2_ increased MMR. This may indicate that negative effects of short term exposures used in our study (i.e. weeks) can be overcome in the long term (i.e. years) (Crespel et al., 2019). As such, future work needs to determine whether interactions between CO_2_ and temperature or O_2_ occur over longer time scales. The effect of CO_2_ on metabolic rate in our study was only identifiable by including the rise in CO_2_ which co-occurs when environmental O_2_ declines. Without the additional data from increased CO_2_ exposures at lower O_2_ levels the effect of ambient CO_2_ on MMR at normoxia was not significant. Although the best model of SMR also included a negative effect of CO_2_ this was predicted to be small with confidence intervals overlapping zero (Figure 1). Additionally, removal of CO_2_ as an explanatory variable from the model of SMR did not greatly impair model fit (ΔAICc < 2) which indicates that the effect of CO_2_ on SMR was not critically important for overall model performance.

As CO_2_ increases in the environment when O_2_ declines the negative effect of CO_2_ has a greater impact on MMR at lower O_2_ levels (Figure 3 C, D, and E). Decreased MMR when fish are exposed to acutely increased CO_2_ is usually thought to be a result of an internal acidosis causing a decrease in Hb-O_2_ binding affinity, reducing the capacity of O_2_ transport in the blood (Heuer & Grosell, 2016). However, fish have well developed acid-base regulatory mechanisms and blood sampling showed that sea bass in normoxia had fully compensated for the effects of increased environmental CO_2_ on blood pH by compensatory accumulation of extra HCO_3_^-^, resulting in no changes in Hb-O_2_ binding affinity (Figure 5 and S1). Additionally, we have recently shown that sea bass are able regulate blood pH when exposed to concurrent progressive hypercapnia during progressive hypoxia over the course of several hours and have higher Hb-O_2_ binding affinity in these conditions when compared to fish exposed to progressive hypoxia with no concurrent hypercapnia (Montgomery et al., 2019). In addition, we did not see significant changes in Hb levels or Hct between CO_2_ treatments (Figure 7). We conclude that the negative effect of increased CO_2_ on MMR is unlikely to be related to changes in O_2_ transport capacity in the blood. Whilst beyond the scope of the present study we can speculate that instead CO_2_ may affect MMR via changes in mitochondrial metabolism (Leo et al., 2017; Strobel et al., 2012, 2013), cardiac performance (Crespel et al., 2019; Perry & Abdallah, 2012) or via shifts from aerobic to anaerobic metabolic pathways (Michaelidis et al., 2007). Indeed, recent results show that CO_2_ impacts mitochondrial function of sea bass from the Atlantic population in combination with acute warming (Howald et al., 2019)

While the effect of CO_2_ on MMR was part of the best supported model it is uncertain whether it would cause biologically relevant impacts. Previous research has linked declines in MMR caused by increased CO_2_ with decreased swimming performance (Lefevre, 2019) but it is unknown if the relatively small changes in MMR shown in the present study would translate to other aspects of whole animal performance. This is especially the case as predictions of aerobic scope from the combined best supported models of MMR and SMR essentially show no effect of CO_2_ on aerobic scope at O_2_ levels >10 kPa (Figure 4 C, D, and E). This occurs because predicted effects of CO_2_ act in the same direction for both SMR and MMR. As changes in aerobic scope typically predict environmental impacts on processes such as growth and reproduction (Clark et al., 2013; Pörtner et al., 2017) we would predict that climate change relevant CO_2_ increases have negligible effects on these endpoints. However, the effect of CO_2_ may have important consequences not reflected in changes in aerobic scope. In our most extreme treatment (22 °C, ∼30 % air saturation, end of century CO_2_) we observed 30 % mortality of sea bass after exhaustive exercise. Fish exercised at the same temperature and O_2_ levels in ambient CO_2_ conditions showed no mortality (and no mortality was observed in any other treatment combinations) – consequently it appears the elevated CO_2_ during hypoxia in a future ocean scenario (which was approximately 1100 µatm higher than in the present day CO_2_ scenario) may impair recovery from exercise when O_2_ is limiting. Mortality in fish post-exercise has been theorised to result from intracellular acidosis generated during anaerobic respiration (Wood et al., 1983). Therefore, the greater increase in CO_2_ during hypoxia in the future ocean CO_2_ scenario may either exacerbate the intracellular acidosis caused by anaerobic activity or impair the ability of fish to process anaerobic end products.

### 3.1. Evidence to support OCLTT?

The OCLTT hypothesis suggests climate change will affect fish because combined effects of reduced O_2_ and increasing CO_2_ will synergistically interact, lowering aerobic scope across its thermal performance curve (Pörtner & Peck, 2010), and that changes in aerobic scope can be used as a single metric for predicting whole animal performance (Pörtner, 2012). Our data provides some support for the OCLTT hypothesis as the observed interactions between increased temperature and reduced O_2_ on aerobic scope would be expected to result in changes to the thermal performance curve as predicted by Pӧrtner and Farrell (2008). However, our highest temperature treatment did not decrease MMR or aerobic scope and so we cannot confirm whether interactive effects between temperature and hypoxia would follow predictions of the OCLTT above the optimum temperature of aerobic scope. In contrast to hypoxia, the effects of CO_2_ did not follow predictions from the OCLTT as CO_2_ did not interact with either temperature or hypoxia and had minimal impacts on aerobic scope.

The interactive effects of O_2_ and temperature on the MMR of sea bass in this study closely resemble predictions of the metabolic niche framework which Ern (2019) proposed as an update to the OCLTT hypothesis. In particular the concept of aerobic scope isopleths (where aerobic scope remains constant across a range of temperatures as a result of changes in O_2_ or vice versa) is supported by our data, showing that aerobic scope of sea bass would be expected to remain constant across an 8 °C temperature range at an O_2_ level of ∼6 kPa (Figure 3 A and B). As such, we would support Ern’s suggestion to experimentally assess how aerobic scope affects important processes, such as growth, independently of changes in temperature and O_2_ by utilising these isopleths. Although increased aerobic scope has long been linked to improved individual fitness there is a still a lack of evidence to confirm this relationship occurs – as such the assumption that peak physiological fitness occurs when aerobic scope peaks (central to the OCLTT) may be erroneous.

Alternatively, Deutsch et al. (2015) have suggested a framework which aims to predict impacts of climate change on the physiological suitability of a habitat for a species via a metabolic index, relating the ratio of O_2_ supply to resting metabolic demand (i.e. SMR), rather than aerobic scope. This metabolic index appears to provide a tool for predicting the biogeographical distributions of species with the biogeographical distribution limits of many marine species corresponding to a metabolic index of ∼2-5 (Deutsch et al., 2015). This approach has been supported by experimental work with black sea bass, Centropristis striata, (Slesinger et al., 2019) and Roman sea bream, Chrysoblephus laticeps, (Duncan et al., 2020) which showed that the metabolic index can accurately predict changes in population distributions of these species . Applying the principle of the metabolic index to our data suggests that temperatures of 22 °C are close to the upper temperature limits of European sea bass from the North-East Atlantic sub-population (metabolic index at 22 °C and 20 kPa O_2_ was ∼5).

## 4. Conclusion

In summary, our research shows that aerobic scope of European sea bass will increase with expected warming in the North-East Atlantic, and that even extreme summer temperatures (∼22 °C) at the end of the century will positively impact on the absolute aerobic performance of sea bass. However, synergistic interactions between warming and reduced O_2_ indicate that hypoxic conditions will have greater impacts on sea bass in future oceans. Increased CO_2_ levels showed no interactions with either temperature or O_2_ changes but were predicted to cause a small decline in MMR – although this had little impact on aerobic scope because increased CO_2_ caused a trend for decreased SMR. Sea bass fully compensated blood pH for increased CO_2_ levels and increases in SMR and reductions in MMR with temperature were not linked to changes in O_2_ transport. Despite end of century CO_2_ levels having minimal effects on aerobic scope, they did cause increased mortality of fish recovering from exercise in the more extreme hypoxic scenario (∼30 % air saturation) at 22 °C. This effect would not have been observed without including expected increases in CO_2_ as O_2_ declines in hypoxia treatments. Thus, environmentally relevant changes in CO_2_ during hypoxia may lead to important threshold effects which could be missed if experiments only consider changes in CO_2_ related to atmospheric concentrations. Interactive effects of temperature and O_2_ support predictions from the oxygen-and temperature-limited metabolic niche framework proposed as an update to the OCLTT hypothesis by Ern (2019), however the effect of CO_2_ did not support predictions of the OCLTT (Pörtner & Farrell, 2008). Changes in the metabolic index proposed as a physiological constraint by Deutsch et al. (2015) suggest that despite increases in MMR and aerobic scope future climate change may result in conditions which will begin to constrain growth and reproduction of sea bass in areas where temperatures increase above 22 °C. However, there is a vital need for increased research to link changes in aerobic scope to population relevant metrics such as growth and reproduction to better assess what environmental impacts on aerobic performance may mean for wider populations.

## 5. Materials & Methods

### 5.1. Animal Collection and Husbandry

We collected juvenile sea bass from estuaries and coastal lagoons on the south Dorset coast and Isle of Wight in June 2017. Fish were held in a marine recirculating aquaculture system (RAS) at the University of Exeter for 332 days before experimental work began (see water chemistry data in Table 1). Sea bass were fed a diet of commercial pellet (Horizon 80, Skretting) at a ration of ∼1-2 % body mass three times per week and supplemented with ∼1 % body mass of chopped mussel (Mytilus edulis) once per week. All experimental procedures were carried out under a UK Home Office licence (P88687E07) and approved by the University of Exeter’s Animal Welfare and Ethical Review Board.

**Table 1:**
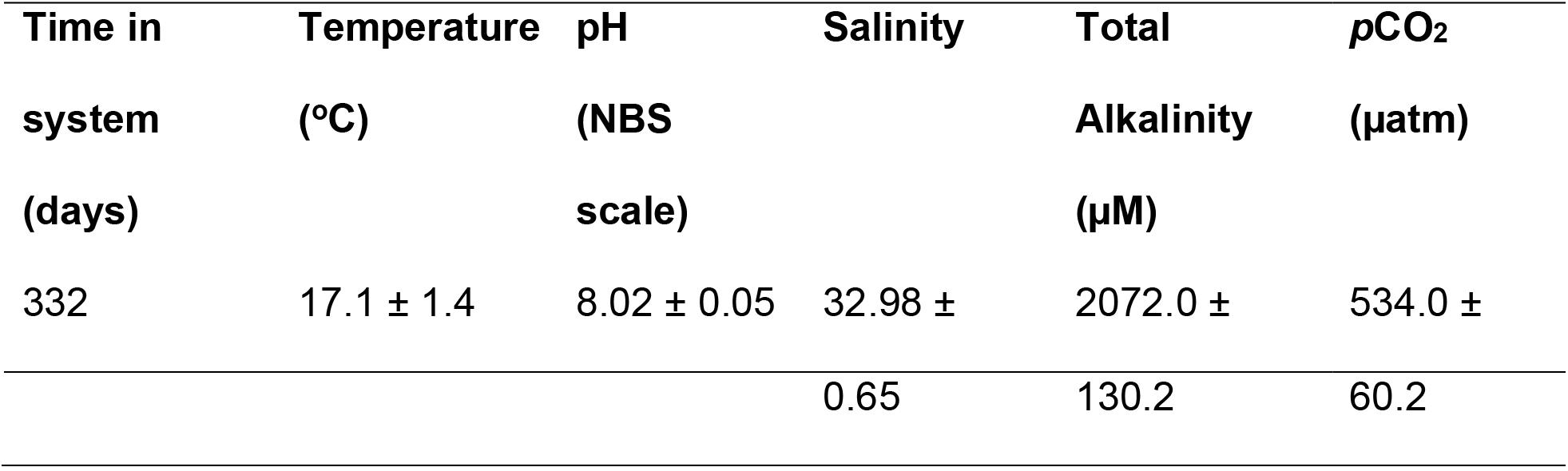
Water chemistry parameters of the recirculating aquaculture system in which sea bass were held prior to experimental work beginning (means ± S.D. shown). Fish were initially held at 15 °C before stock systems were raised to 18 °C approximately 6 months prior to experimental work beginning. The temperature shown in the table represents the mean temperature for the entire time fish were held in the RAS prior to experimental trials beginning.

### 5.2. Treatment conditions

Sea bass were transferred to an experimental RAS, in a temperature-controlled room for a minimum of 14 days acclimation to treatment conditions (Figure 6). Six treatment conditions were used combining three temperatures and two CO_2_ levels in a three x two factorial design (Table 2). Temperature treatments (14, 18 and 22 °C) were chosen to reflect temperature ranges in coastal UK waters from spring to autumn as well as potential future summer temperatures at the end of the century (IPCC, 2014; Tinker et al., 2020). CO_2_ treatments (∼400 & ∼1000 µatm) were chosen to reflect annual average ambient atmospheric CO_2_ levels currently and possible end-of-century ambient atmospheric levels according to an RCP 8.5 scenario (IPCC, 2014). Sea bass were transferred to the experimental RAS at 18 °C before temperatures were adjusted at a rate of 2 °C per day to reach treatment conditions (i.e. 14 or 22 °C). A header tank (∼500 L) in the experimental RAS was used to adjust CO_2_ to the desired level for each treatment before entering the treatment tank which contained the sea bass. For present day CO_2_ treatments (∼400 µatm) the header tank was aerated using CO_2_ scrubbed air to remove excess CO_2_ added to the RAS by the biological filters. For end of century CO_2_ treatments (∼1000 µatm) an Aqua Medic pH computer was used to adjust RAS water to an appropriate pH (7.8). Additionally, treatment tanks were aerated with a gas mix with the appropriate CO_2_ content for each treatment.

**Table 2:** Mean ± S.D. of water chemistry parameters in treatment tanks during treatments at 3 temperature (14, 18, and 22 °C) and 2 CO_2_ levels (∼400 and ∼1000 µatm). Treatment order represents time course of treatments (i.e. treatment 1 was conducted first and treatment 6 last). Total Alkalinity (TA) varied somewhat over time as a result of biological activity and so was adjusted periodically by addition of 1.0 M NaHCO_3_ to restore TA levels back to >2000 µM.

| Parameter | Treatment |  |  |  |  |  |
| --- | --- | --- | --- | --- | --- | --- |
|  | 1 | 2 | 3 | 4 | 5 | 6 |
| Temperature | 17.9 ± | 22.0 ± | 13.9 ± 0.1 | 21.7 ± 0.5 | 13.9 ± 0.1 | 18.0 ± |
| (°C) | 0.0 | 0.0 |  |  |  | 0.1 |
| $p\text{CO}_2$ ($\mu\text{atm}$) | 460 ± 34 | 375 ± 28 | 333 ± 38 | 1065 ± 172 | 1057 ± 65 | 973 ± |
|  |  |  |  |  |  | 114 |
| pH (NBS) | 8.06 ± | 8.12 ± | 8.14 ± 0.04 | 7.79 ± 0.03 | 7.81 ± 0.04 | 7.83 ± |
|  | 0.03 | 0.02 |  |  |  | 0.05 |
| Salinity | 32.9 ± | 32.5 ± | 33.3 ± 0.3 | 32.6 ± 0.3 | 34.6 ± 0.5 | 33.8 ± |
|  | 0.6 | 2.9 |  |  |  | 0.6 |
| TA ($\mu\text{M}$) | 1935 ± | 1745 ± | 1861 ± 88 | 2147 ± 292 | 2393 ± 188 | 2170 ± |
|  | 93 | 26 |  |  |  | 20 |

Measurements of treatment tank pH (NBS scale), salinity, temperature, and a 12 mL water sample to measure Dissolved Inorganic Carbon (DIC) were taken every 2-3 days. Seawater DIC analysis was conducted using a custom built system described in detail by Lewis et al. (2013). Data for pH, salinity, temperature and DIC were then input into the seawater carbon calculator programme, CO2SYS (Pierrot et al., 2006) to calculate pCO_2_ based on the equilibration constants refitted by Dickson and Millero (1987), and KSO_4_ dissociation constants from Dickson (1990) (Table 2).

### 5.3. Respirometry measurements (SMR, MMR, and O_2crit_)

Rates of oxygen consumption (ṀO_2_) were made as a proxy of metabolic rate using an intermittent-flow respirometer system, details of which can be found in Montgomery et al. (2019), set-up following recommendations by Svendsen et al. (2016). Sea bass were starved for 72 hours prior to the start of measurements to ensure that metabolism was not affected by the specific dynamic action of digestion (Chabot et al., 2016). Individual sea bass were then transferred to respirometer chambers and left to acclimate for a minimum of 12 hours overnight before measurements of ṀO_2_ began. For each treatment all respirometry measurements were conducted in two groups (hereafter referred to as respirometry group), with five fish being measured simultaneously for each group. Following the 12 hour acclimation period we measured ṀO_2_ of each sea bass for ∼3-4 hours (from ∼6 am to ∼10 am) before hypoxia tolerance was assessed using a critical O_2_ tension (O_2crit_) trial (Figure 6), following protocols set out in Montgomery et al. (2019). Carbon dioxide levels in the water were simultaneously increased as O_2_ declined during O_2crit_ trials to reflect the natural rise in CO_2_ during hypoxic events in aquatic systems (Melzner et al., 2013; Montgomery et al., 2019). During O_2crit_ trials water pH, temperature, salinity, and DIC were measured every hour to calculate water carbonate chemistry. Changes in system pCO_2_ and pH during O_2crit_ trials for each treatment are given in supplementary materials (Table S1).

O_2crit_ trials were stopped once a minimum of three consecutive ṀO_2_ measurements showed a transition from an oxy-regulating to oxy-conforming state for each fish. Following completion of O_2crit_ trials the respirometer system was aerated with ambient air (CO_2_ ∼400 µatm) or a 0.1 % CO_2_ in air gas mix (CO_2_ ∼1000 µatm) to rapidly restore O_2_ levels to normoxia and CO_2_ to the appropriate treatment level. Sea bass were left to recover in respirometers, for a minimum of one hour post-trial, until O_2_ levels reached ∼ 21 kPa O_2_ (∼100 % air saturation) before removing the fish and measuring background respiration for a minimum of one hour (six measurement cycles) for all respirometers immediately post trial. Each sea bass was then placed in an individual ∼10 L isolation tank which was subsequently fed by the respirometry system sump (at a rate of ∼4 L min^-1^) to maintain treatment conditions (with overflowing water from the isolation tanks recirculating back to the sump).

After fish had rested overnight in isolation tanks, MMR was measured for each fish (using an exhaustive chase protocol; Norin and Clark, 2016) on three consecutive days (with overnight recovery in between) at three different levels of O_2_ (100, 60 and 30 % air saturation) with increasing CO_2_ levels for each O_2_ level as detailed for O_2crit_ trials (Figure 6). The appropriate O_2_ and CO_2_ level was achieved by aerating isolation boxes with a mix of N_2_, O_2_ and CO_2_ (G400 Gas mixing system, Qubit Biology Inc.) at a rate of 5 L min^-1^. Fish were exposed to the new O_2_ and CO_2_ conditions for ∼2 hours before chase protocols were conducted. Chase protocols and subsequent respirometry measurements were conducted at the appropriate temperature, O_2_, and CO_2_ conditions for each treatment. Measurements of water pH, temperature, salinity, and DIC were taken for each isolation tank, the chase tank (after all fish were chased) and the respirometer system (during ṀO_2_ measurements) to calculate water carbonate chemistry. Temperature, O_2_, and CO_2_ conditions for all MMR trials are given in Table S2.

For all ṀO_2_ measurements dissolved O_2_ concentration (% air saturation) was measured continuously (frequency ∼1 Hz) in respirometer chambers using a fibre optic O_2_ optode mounted in the recirculation loop of the respirometer chamber.

These optodes were linked to two Firesting Optical Oxygen Meters (Pyro Science, Aachen, Germany) which were connected to a PC running AquaResp 3 software which automatically logged all measurements.

ṀO_2_ was automatically calculated by the AquaResp3 software by fitting a linear regression to the O_2_ versus time data for each measurement period. The slope (s) of this regression (kPa O_2_ h^-1^) was then used to calculate ṀO_2_ (mg O_2_ kg^-1^ h^-1^) using the equation outlined by Svendsen et al.(2016):

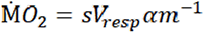

where V_resp_ is the respirometer volume minus the volume of the fish (L), α is the solubility of O_2_ in water (mgO_2_ L^-1^ kPa^-1^) for the relevant salinity and temperature, and m is the mass of the fish (kg). For the purpose of establishing the impacts of reduced O_2_ on MMR and determining O_2crit_ values, the O_2_ level of each measurement period was defined as the mean dissolved O_2_ measurement over the measurement period. The mean background respiration for each respirometer over the 1 hour post-trial measuring period (average was < 2 % of fish ṀO_2_) was subtracted from ṀO_2_ measurements. Background corrected ṀO_2_ was then scaled to an average individual mass of 120 g using a mass exponent of 0.89 prior to subsequent analysis (Jerde et al., 2019).

We calculated SMR of each fish as the mean of the lowest 10 ṀO_2_ measurements from the ∼3-4 hour period prior to O_2crit_ trials in which mean dissolved O_2_ saturation was >80 % air saturation. The critical O_2_ tension (O_2crit_) of each individual fish was then calculated using ṀO_2_ measurements from O_2crit_ trials with function ‘calcO2crit’ from package ‘fishMO2’ (Chabot et al. 2016) in R v.3.6.3 (R Core Team, 2020), using the estimated SMR of each individual, as detailed in the supplementary material of Claireaux & Chabot (2016). Finally, we defined MMR as the single highest measurement of ṀO_2_ in the one hour period immediately following exercise to exhaustion (Norin & Clark, 2016). This point usually occurred during the first measurement period immediately after each fish was moved to the respirometer chamber i.e. ∼ 2-5 minutes after the cessation of the chase protocol. However, for some fish in normoxia spontaneous activity inside the respirometer chamber during SMR or O_2crit_ trials resulted in instantaneous measurements of ṀO_2_ higher than those noted following chase protocols. In these occasions this higher value of ṀO_2_ was used as the estimated MMR for that fish (n = 2 out of 65 fish).

### 5.4. Blood chemistry and Hb-O_2_ affinity measurements

Following MMR measurements, sea bass were left overnight in the isolation boxes before blood samples were taken (Figure 6), following methods outlined in Montgomery et al. (2019), from each fish in normoxic conditions and at the relevant treatment temperature and CO_2_ level (Table S3). We then measured extracellular pH (pH_e_), haematocrit (Hct), TCO_2_, haemoglobin content (Hb), plasma glucose, and plasma lactate and calculated pCO_2_ and HCO_3_^-^ following methods detailed in Montgomery et al. (2019). We also followed the freeze-and-thaw method to measure intracellular pH of RBCs (pH_i_) as described by Zeidler and Kim (1977), and validated by Baker et al. (2009). All measurements or storage of blood for subsequent analysis occurred within 10 minutes of blood sampling. Finally, we measured Hb-O_2_ affinity using a Blood Oxygen Binding System (BOBS, Loligo systems), detailed in general in Oellermann et al. (2014) and specifically for fish blood in Montgomery et al. (2019).

**Figure 6:**
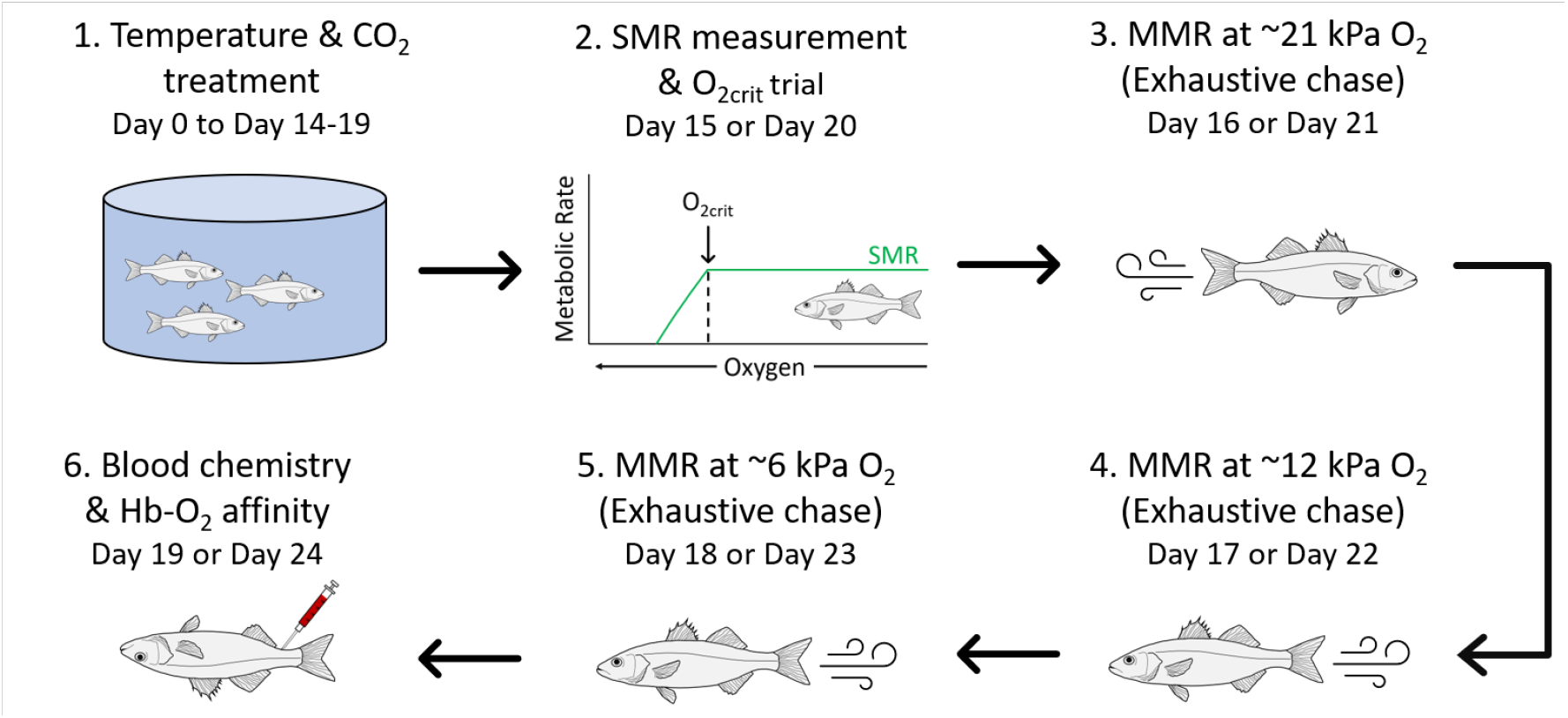
Summary of the timeline over which experimental end points were measured. Sea bass were acclimated for a minimum of 14 days to a temperature (14, 18, 22 °C) and CO_2_ (∼400 µatm or ∼1000 µatm ambient CO_2_) treatment before measurements of SMR/O_2crit_, MMR (at ∼21 kPa, ∼12 kPa, and ∼6 kPa O_2_), and blood chemistry/Hb-O_2_ affinity were obtained on consecutive days for each individual.

### 5.5. Statistical Analysis

#### 5.5.1. Respirometry data analysis

We conducted all statistical analysis in R v3.6.3 (R Core Team, 2020). Results are reported as mean ± S.E unless otherwise stated. Sample sizes for respirometry data can be seen in Table S4. The effects of temperature, O_2_, and CO_2_ on individual physiological performance metrics (SMR, MMR, and O_2crit_) were analysed using separate general linear mixed-effects models (GLMM) in package ‘lme4’ (Bates et al., 2015; Pinheiro et al., 2018). All models included respirometry group as a random intercept term to account for potential tank effects introduced during respirometry measurements. For each physiological metric the best supported model was determined as the model with the lowest corrected Akaike’s information criterion, AICc (Burnham & Anderson, 1998; Hurvich & Tsai, 1989). Residual diagnostic plots of each GLMM were then assessed using package ‘DHARMa’ to confirm validity of model fit (Hartig, 2020). Once the best supported model for each physiological parameter was identified (see Table S5 for model comparisons) predictions were made across a range of temperatures, O_2_ levels, and CO_2_ levels to visualise combined effects of these variables on the physiology of seabass. We then used function bootMer from lme4 (Pinheiro et al., 2018) to calculate 95 % confidence intervals of model predictions.

#### 5.5.2. Blood chemistry and Hb-O_2_ affinity data analysis

Measurements of blood chemistry parameters (pH_e_, pH_i_, pCO_2_, HCO_3_^-^, P_50_, Hills’ number, Hct, Hb, lactate, and glucose) were analysed using the ambient water temperature of the treatment and using a categorical CO_2_ level of low (i.e. ∼400 µatm treatment) or high (i.e. ∼1000 µatm treatment). Measurements were analysed using a type III sum of squares two-way ANOVA (to account for unequal sample sizes). Post hoc-tests were then conducted on least-square means generated by package ‘emmeans’ (Lenth, 2020), with Tukey adjusted p-values for multiple comparisons. Model residuals from analysis of plasma lactate and glucose measurements did not meet the assumptions of normality or equal variances required by two-way ANOVA, as such this data was analysed using the alternative non-parametric Kruskal-wallis test.

Blood-oxygen binding parameters (P_50_ and Hills’ number) could not be obtained for fish in the 14 °C and high CO_2_ treatment as a result of an equipment failure. As such these data were analysed using a one-way ANOVA. If statistical assumptions of one-way ANOVA were not met then data were analysed using the non-parametric Kruskal-Wallis test. Sample sizes for blood chemistry data can be seen in Table S4.

## Acknowledgements

This work was supported by a NERC GW4+ Doctoral Training Partnership studentship from the Natural Environment Research Council [NE/L002434/1] with additional funding from CASE partner, The Centre of Fisheries and Aquaculture Science (Cefas) to D.W.M., and from the Biotechnology and Biological Sciences Research Council (BB/D005108/1 and BB/J00913X/1) and NERC (NE/H017402/1) to R.W.W. The authors would like to thank the Marine Management Organisation and Natural England for granting permits (Marine Management Organisation permit #030/17 & Natural England permit #OLD1002654) to collect wild sea bass for use in this study, and Simon Pengelly and the Southern Inshore Fisheries and Conservation Authority for assistance with fish collections. We would also like to thank the aquarium staff, particularly Steven Cooper, Alice Walpole and Rebecca Turner, of the Aquatic Resource Centre at the University of Exeter for assistance with fish husbandry and system water chemistry, and finally Dr Cosima Porteus for assistance with blood chemistry measurements.

## Data availability

Data will be made publicly available via the University of Exeter’s online repository if the manuscript is accepted: https://ore.exeter.ac.uk/repository/

## Supplementary materials

**Table S1:** Measurements of pCO_2_ (µatm, top row) and pH (NBS scale, bottom row) as O_2_ declined during O_2crit_ trials for each treatment. All values are mean ± S.D. apart from for Treatment 3 at ∼30% dissolved O_2_ where only one measurement of pCO_2_ and pH was made and calculations of mean and S.D. were not possible

| ~Dissolved $\text{O}_2$<br>(% air saturation) | Treatment | | | | | |
| --- | --- | --- | --- | --- | --- | --- |
|  | 14 °C | 18 °C | 22 °C | 14 °C | 18 °C | 22 °C |
| | Present $\text{CO}_2$ | Present $\text{CO}_2$ | Present $\text{CO}_2$ | Future $\text{CO}_2$ | Future $\text{CO}_2$ | Future $\text{CO}_2$ |
| 100 | 377 ± 23 | 483 ± 20 | 455 ± 35 | 1019 ± 50 | 1070 ± 10 | 1172 ± 89 |
| | 8.14 ± 0.02 | 8.09 ± 0.02 | 8.10 ± 0.04 | 7.79 ± 0.01 | 7.78 ± 0.00 | 7.80 ± 0.03 |
| 80 | 418 ± 9 | 508 ± 10 | 486 ± 47 | 1149 ± 2 | 1156 ± 86 | 1209 ± 18 |
| | 8.10 ± 0.01 | 8.07 ± 0.01 | 8.07 ± 0.04 | 7.74 ± 0.01 | 7.75 ± 0.02 | 7.79 ± 0.02 |
| 60 | 477 ± 25 | 613 ± 90 | 497 ± 23 | 1383 ± 125 | 1315 ± 73 | 1335 ± 92 |
| | 8.05 ± 0.02 | 7.99 ± 0.01 | 8.07 ± 0.02 | 7.67 ± 0.02 | 7.69 ± 0.02 | 7.75 ± 0.03 |
| 40 | 625 ± 44 | 746 ± 40 | 577 ± 46 | 1637 ± 83 | 1488 ± 24 | 1658 ± 237 |
| | 7.94 ± 0.03 | 7.92 ± 0.02 | 8.01 ± 0.04 | 7.59 ± 0.01 | 7.64 ± 0.00 | 7.66 ± 0.05 |
| 30 | 822 | 1104 ± 155 | 647 ± 18 | 1973 ± 159 | 1964 ± 134 | 2179 ± 63 |
| | 7.83 | 7.76 ± 0.06 | 7.96 ± 0.01 | 7.51 ± 0.04 | 7.53 ± 0.02 | 7.55 ± 0.01 |
| 20 | 1162 ± 2 | 2103 ± 79 | 808 ± 35 | 2491 ± 238 | 2416 ± 265 | 2547 ± 266 |
| | 7.69 ± 0.00 | 7.50 ± 0.02 | 7.88 ± 0.02 | 7.42 ± 0.05 | 7.45 ± 0.03 | 7.48 ± 0.06 |
| 15 | 1734 ± 92 | 2431 ± 663 | 1318 ± 108 | 3099 ± 448 | 2579 ± 4 | 3380 ± 354 |
| | 7.53 ± 0.02 | 7.45 ± 0.11 | 7.68 ± 0.04 | 7.32 ± 0.07 | 7.41 ± 0.00 | 7.36 ± 0.06 |

**Table S2:**
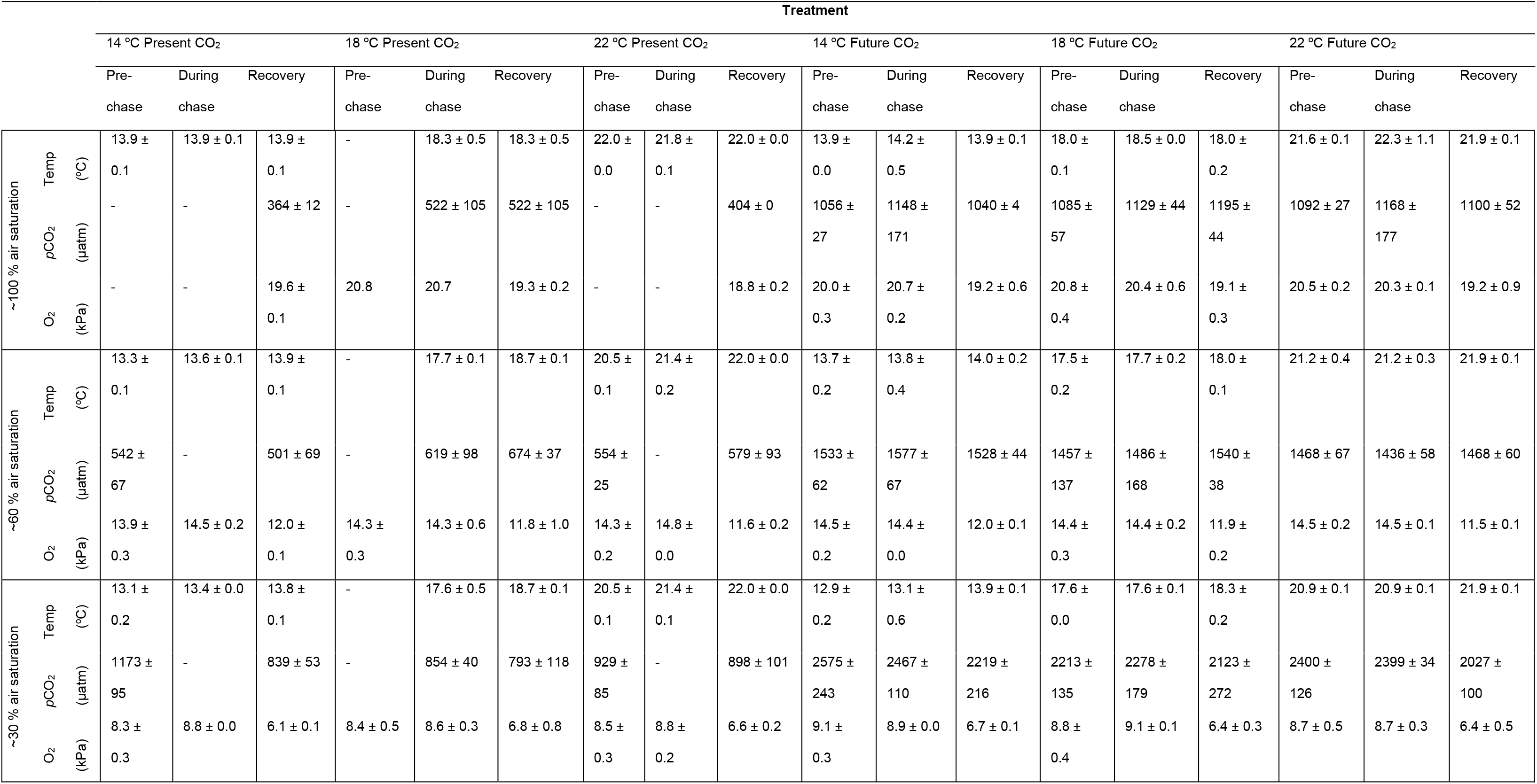
_Temperature (_°_C), pCO_2 _(µatm), and O_2 _(kPa) conditions sea bass were exposed to pre-chase, during chase, and when recovering from exercise during measurements of MMR. All values are presented as mean ± S.E. - indicates measurements for which data was not collected. For the 14_ °_C present day CO_2 _treatment missing pre-chase values would be expected to be extremely similar to values recorded during chase as shown by values for all other treatments._

**Table S3:** Temperature and pCO_2_ levels fish were exposed to when they were anaesthetised immediately prior to blood sampling. All measures are given as mean ± S.D.

| Parameter | Treatment |  |  |  |  |  |
| --- | --- | --- | --- | --- | --- | --- |
|  | 14 °C | 18 °C | 22 °C | 14 °C | 18 °C | 22 °C |
|  | Present CO<sub>2</sub> | Present CO<sub>2</sub> | Present CO<sub>2</sub> | Future CO<sub>2</sub> | Future CO<sub>2</sub> | Future CO<sub>2</sub> |
| Temperature (°C) | 13.1 ± 0.1 | 17.4 ± 0.1 | 21.2 ± 0.3 | 13.3 ± 0.3 | 17.8 ± 0.1 | 21.3 ± 0.2 |
| $p\text{CO}_2$ ($\mu\text{atm}$) | 350 ± 13 | 445 ± 52 | 398 ± 28 | 1130 ± 33 | 1103 ± 47 | 1221 ± 35 |

**Table S4:** Samples sizes for each measurement in all treatments. We were unable to make measurements of P_50_ or Hills number for the 14 °C future CO_2_ scenario due to an equipment failure.

| Measurement | Treatment |  |  |  |  |  |
| --- | --- | --- | --- | --- | --- | --- |
|  | 14 °C | 18 °C | 22 °C | 14 °C | 18 °C | 22 °C |
|  | Present CO<sub>2</sub> | Present CO<sub>2</sub> | Present CO<sub>2</sub> | Future CO<sub>2</sub> | Future CO<sub>2</sub> | Future CO<sub>2</sub> |
| SMR | 10 | 10 | 10 | 10 | 10 | 10 |
| O<sub>2crit</sub> | 10 | 10 | 10 | 10 | 10 | 10 |
| MMR at ~21 kPa O<sub>2</sub> | 9 | 10 | 10 | 12 | 10 | 10 |
| MMR at ~12 kPa O<sub>2</sub> | 10 | 7 | 10 | 10 | 10 | 10 |
| MMR at ~6 kPa O<sub>2</sub> | 10 | 7 | 10 | 10 | 10 | 10 |
| Blood pH<sub>e</sub> | 10 | 6 | 9 | 10 | 9 | 7 |
| Plasma $p\text{CO}_2$ | 10 | 6 | 9 | 10 | 9 | 7 |
| Plasma HCO<sub>3</sub><sup>-</sup> | 10 | 6 | 9 | 10 | 9 | 7 |
| RBC pH<sub>i</sub> | 10 | 6 | 8 | 9 | 9 | 7 |
| P<sub>50</sub> | 10 | 6 | 9 | NA | 8 | 7 |
| Hills number | 10 | 6 | 9 | NA | 8 | 7 |
| Haematocrit | 10 | 6 | 9 | 10 | 9 | 7 |
| Haemoglobin | 7 | 6 | 9 | 10 | 9 | 7 |
| Plasma lactate | 9 | 6 | 9 | 7 | 8 | 7 |
| Plasma glucose | 9 | 6 | 10 | 7 | 8 | 7 |

**Table S5:**
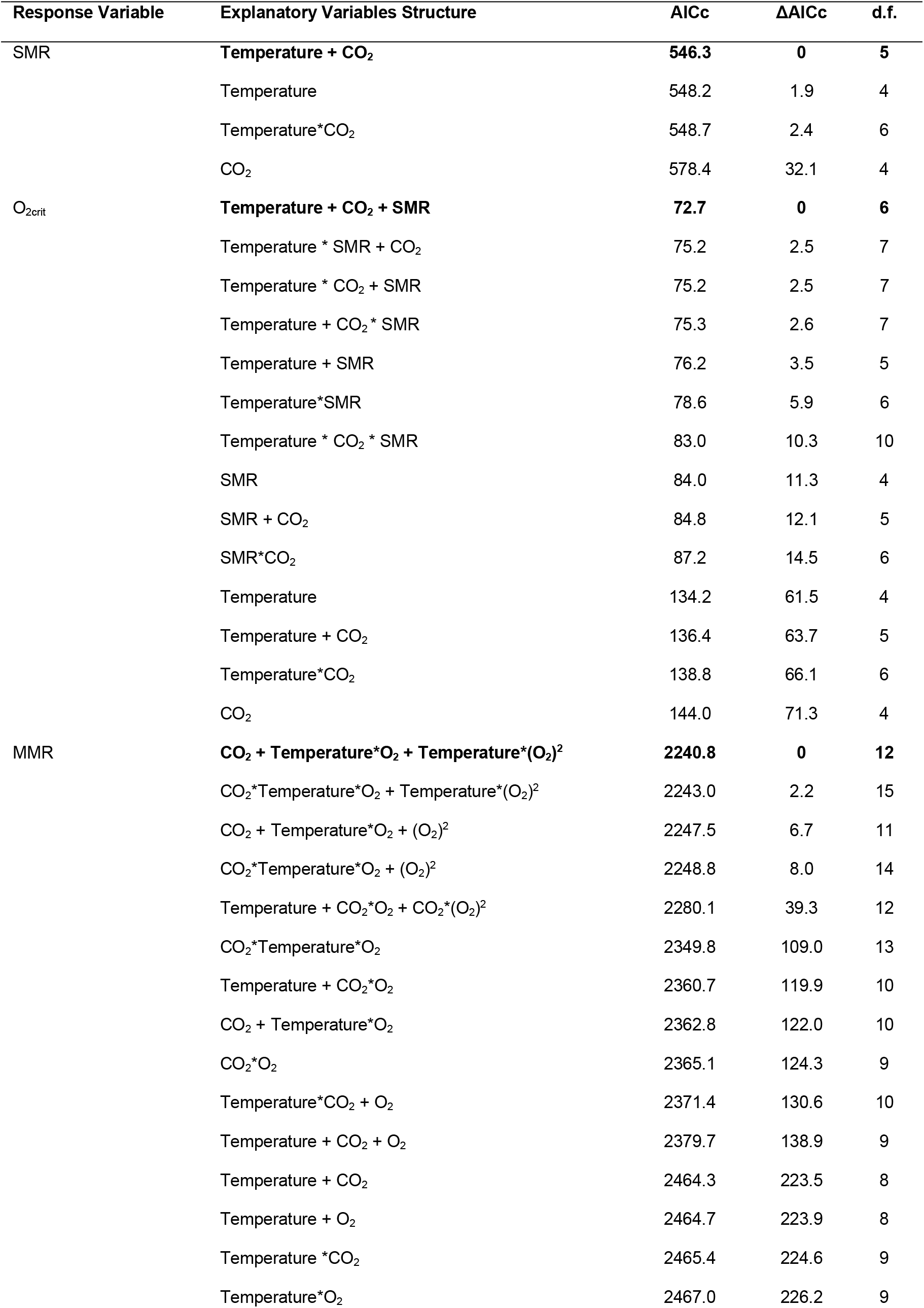
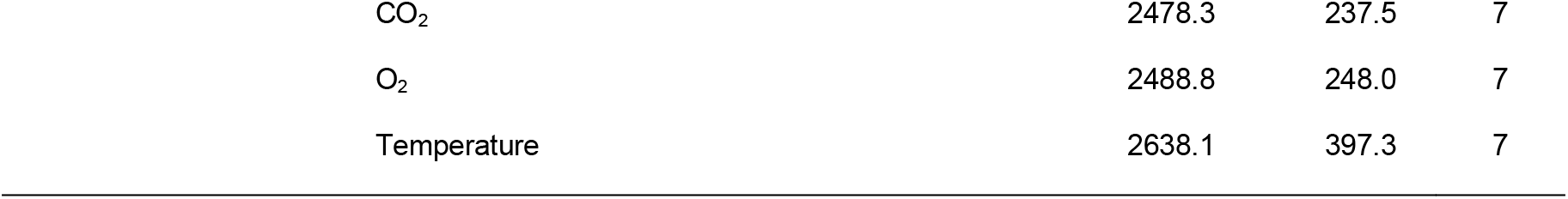
Structure of models and model comparison results explaining variation in SMR, O_2crit_, and MMR of sea bass exposed to differing combinations of temperature, O_2_, and CO_2_. The best supported model for each response variable is highlighted in bold.

**Table S6:** General linear mixed-effects model outputs for analysis of standard metabolic rate. The best supported model was fitted using a Gaussian distribution and included the parameters temperature and CO_2_ as explanatory variables and group ID as a random intercept term. Parameter effects are compared against a reference level where temperature and CO_2_ are 0. Marginal R^2^ = 0.70, Condition R^2^ = 0.70.Confidence intervals for each parameter were determined from function confint in package lme4. Marginal and conditional R^2^ of the model were determined using function r.squaredGLMM from package MuMIn.

| Parameter | Estimate | Standard Error | Confidence Interval (95%) | t-value | Variance | Standard Deviation |
| --- | --- | --- | --- | --- | --- | --- |
| Best supported model <- lmer(SMR ~Temperature + CO<sub>2</sub> + (1\|Group)) |  |  |  |  |  |  |
| Intercept | -14.77 | 11.62 | -37.35 – 7.81 | -1.27 |  |  |
| Temperature | 6.66 | 0.57 | 5.55 – 7.78 | 11.61 |  |  |
| CO<sub>2</sub> | -0.012 | 0.006 | -0.024 – -0.001 | -2.04 |  |  |
| Group |  |  |  |  | 0.0 | 0.0 |

**Table S7:**
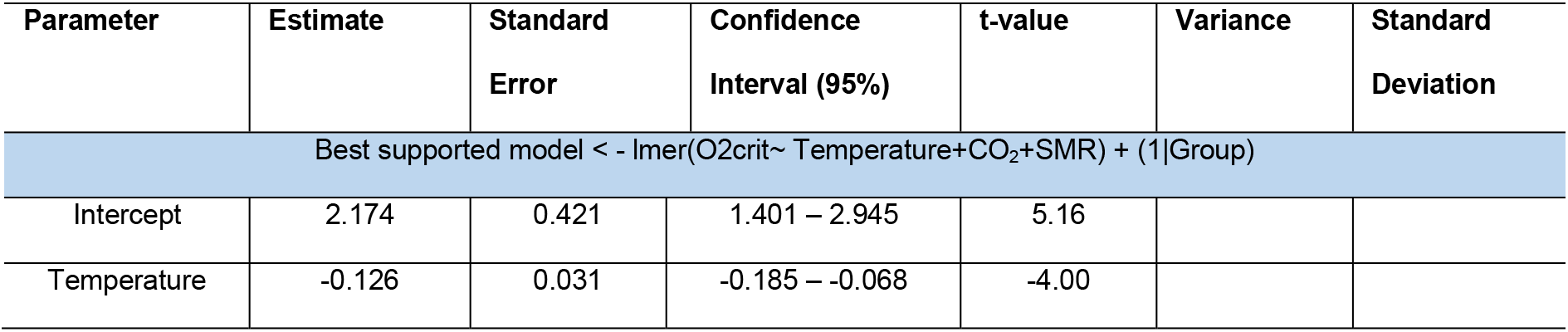
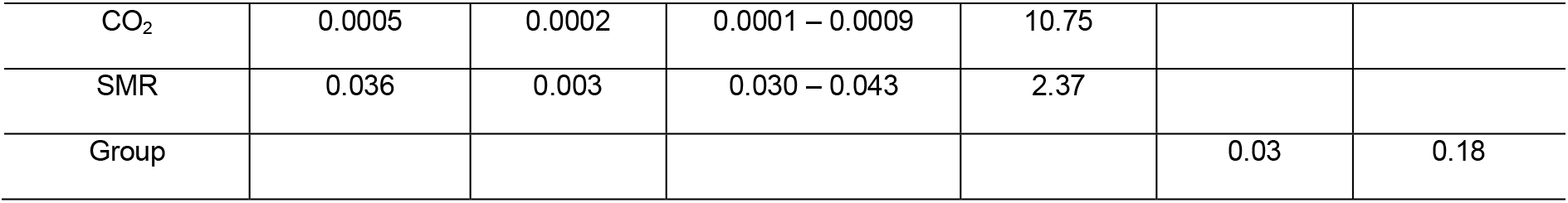
General linear mixed-effects model outputs for analysis of O_2crit_. The best supported model was fitted using a Gaussian distribution and included the parameters temperature, CO_2_, and SMR as explanatory variables and group ID as a random intercept term. Parameter effects are compared against a reference level where temperature, CO_2_, and SMR are 0. Marginal R^2^ = 0.72, Condition R^2^ = 0.77. Confidence intervals for each parameter were determined from function confint in package lme4. Marginal and conditional R^2^ of the model were determined using function r.squaredGLMM from package MuMIn.

**Table S8:** General linear mixed model outputs for analysis of maximum metabolic rate. The best supported model was fitted using a Gaussian distribution and included the parameters temperature and CO_2_ as explanatory variables and group ID as a random intercept term. Parameter effects are compared against a reference level where temperature and CO_2_ are 0. Marginal R^2^ = 0.91, Condition R^2^ = 0.96.Confidence intervals for each parameter were determined from function confint in package lme4. Marginal and conditional R^2^ of the model were determined using function r.squaredGLMM from package MuMIn.

| Parameter | Estimate | Standard Error | Confidence Interval (95%) | t-value | Variance | Standard Deviation |
| --- | --- | --- | --- | --- | --- | --- |
| Best supported model < - lmer(MMR ~ scale(Temperature)*scale(O<sub>2</sub>) + scale(Temperature)*scale(O<sub>2</sub><sup>2</sup>) + scale(CO<sub>2</sub>) + (O<sub>2</sub>\|Fish ID) + (1\|Group) |  |  |  |  |  |  |
| Intercept | 231.99 | 2.58 | 227.01 – 236.98 | 89.96 |  |  |
| scale(Temperature) | 32.53 | 2.55 | 27.60 – 37.46 | 12.76 |  |  |
| scale(O<sub>2</sub>) | 240.35 | 9.50 | 221.92 – 258.74 | 25.31 |  |  |
| scale(Temperature*O<sub>2</sub>) | 41.95 | 8.60 | 25.26 – 58.62 | 4.88 |  |  |
| scale(O<sub>2</sub><sup>2</sup>) | -157.69 | 8.68 | -174.50 – -140.85 | -18.17 |  |  |
| scale(Temperature*O<sub>2</sub><sup>2</sup>) | -25.10 | 8.36 | -41.31 – -8.86 | -3.00 |  |  |
| scale(CO<sub>2</sub>) | -7.52 | 2.58 | -12.08 – -2.97 | -3.20 |  |  |
| Fish ID (Intercept) |  |  |  |  | 90.02 | 9.49 |
| Fish ID (O<sub>2</sub>) |  |  |  |  | 6.53 | 2.56 |
| Group |  |  |  |  | 4.78 x 10<sup>-6</sup> | 0.00 |

**Figure S1:**
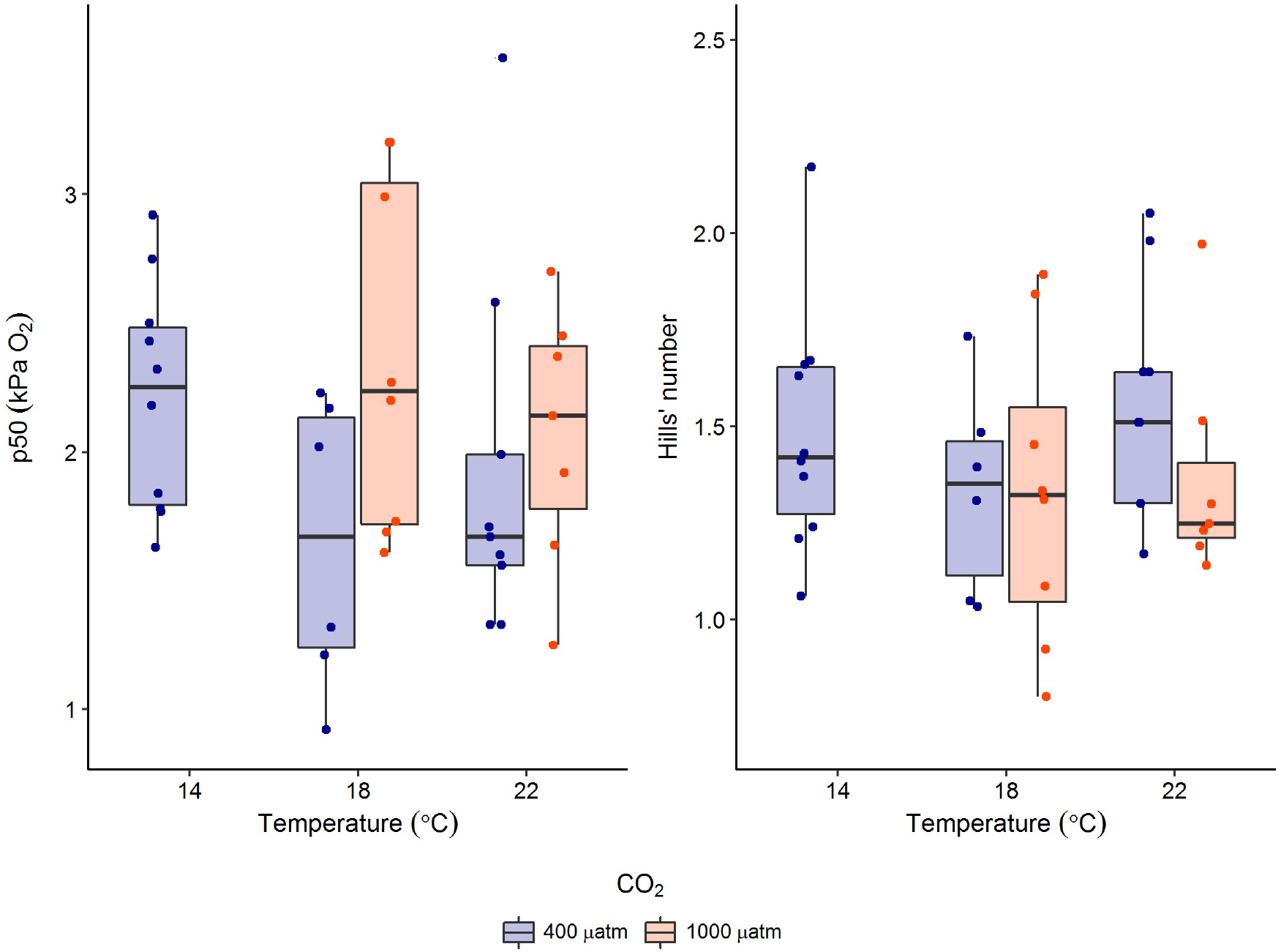
No impacts of temperature and CO_2_ were observed for measurements of haemoglobin-O_2_ binding affinity (measured using P_50_) and Hills’ number. Due to an equipment failure no measurements were possible during the original experimental period for fish at 14 °C exposed to ∼1000 µatm CO_2_.

**Figure S2:**
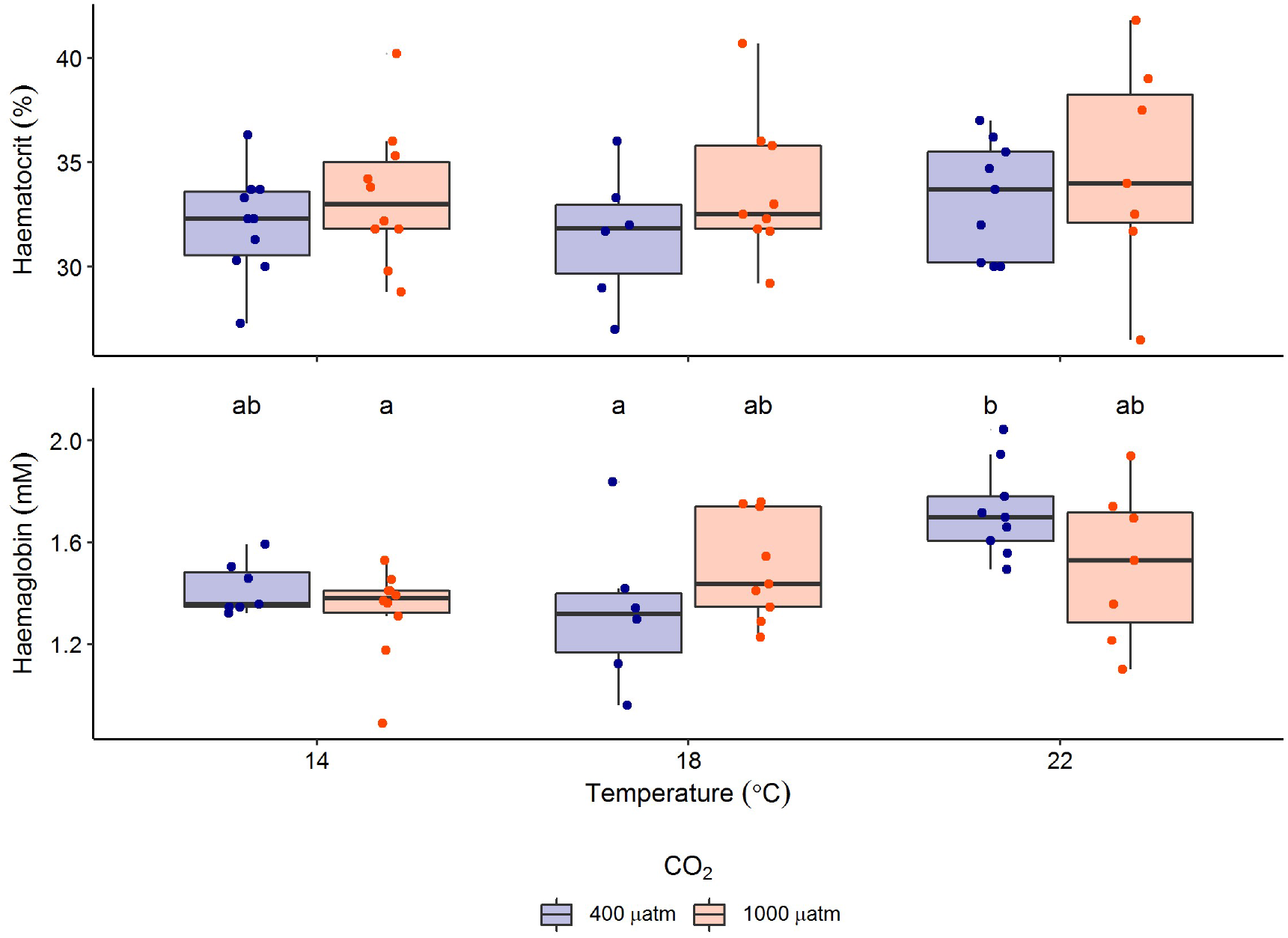
Impact of temperature and CO_2_ on haematological parameters of sea bass. No significant difference in haematocrit were observed between any treatments. Significant difference in haemoglobin content were noted between fish sampled at different temperature and CO_2_ treatments and are represented by different lower case letters.

